# Estimating the functional impact of INDELs in transcription factor binding sites: a genome-wide landscape

**DOI:** 10.1101/057604

**Authors:** Esben Eickhardt, Thomas Damm Als, Jakob Grove, Anders Dupont Boerglum, Francesco Lescai

## Abstract

**Background:** Variants in transcription factor binding sites (TFBSs) may have important regulatory effects, as they have the potential to alter transcription factor (TF) binding affinities and thereby affecting gene expression. With recent advances in sequencing technologies the number of variants identified in TFBSs has increased, hence understanding their role is of significant interest when interpreting next generation sequencing data. Current methods have two major limitations: they are limited to predicting the functional impact of single nucleotide variants (SNVs) and often rely on additional experimental data, laborious and expensive to acquire. We propose a purely bioinformatic method that addresses these two limitations while providing comparable results.

**Results:** Our method uses position weight matrices and a sliding window approach, in order to account for the sequence context of variants, and scores the consequences of both SNVs and INDELs in TFBSs. We tested the accuracy of our method in two different ways. Firstly, we compared it to a recent method based on DNase I hypersensitive sites sequencing (DHS-seq) data designed to predict the effects of SNVs: we found a significant correlation of our score both with their DHS-seq data and their prediction model. Secondly, we called INDELs on publicly available DHS-seq data from ENCODE, and found our score to represent well the experimental data. We concluded that our method is reliable and we used it to describe the landscape of variation in TFBSs in the human genome, by scoring all variants in the 1000 Genomes Project Phase 3. Surprisingly, we found that most insertions have neutral effects on binding sites, while deletions, as expected, were found to have the most severe TFBS-scores. We identified four categories of variants based on their TFBS-scores and tested them for enrichment of variants classified as pathogenic, benign and protective in ClinVar: we found that the variants with the most negative TFBS-scores have the most significant enrichment for pathogenic variants.

**Conclusions:** Our method addresses key shortcomings of currently available bioinformatic tools in predicting the effects of INDELs in TFBSs, and provides an unprecedented window into the genome-wide landscape of INDELs, their predicted influences on TF binding, and potential relevance for human diseases. We thus offer an additional tool to help prioritising non-coding variants in sequencing studies.

## Background

The binding of transcription factors (TFs) to sequence-specific binding sites is crucial for directing gene expression in a time-dependent and cell-selective manner [1,2], and variants in TF binding sites (TFBSs) are thus likely to have important regulatory effects. Variants in transcription factor binding sites have already been associated to various diseases [3,4,5] including diabetes, [6] retinitis pigmentosa [7] and cancers [8,9]. It has become increasingly clear that disease- and trait-associated variants are enriched in regions of TF-binding [10,11], and therefore the need for methods that can estimate their functional impact has grown. The functional importance of TFBSs is supported by their conservation in evolution and the lower levels of variation across binding sites compared to their flanking regions [12,13]. Studies have also shown that the binding of TFs to TFBSs has important roles in chromatin remodelling [14].

The increased availability of next-generation sequencing (NGS) data increases the number of variants discovered in TFBSs: understanding the role of these novel and rare variants is fundamental to the analysis and interpretation of NGS data. However, while it is relatively straightforward to predict the biological effects of variants in the protein-coding regions, it is much harder to interpret their effects when they are positioned in non-coding regions. Available methods depend on multiple sources of data, expensive and laborious to acquire (e.g. data on DNase I sensitivity or histone modifications), and yet are limited to predicting the effects of single nucleotide variants SNVs [15–17].

One method that is often used to predict the probability of a TF binding to a specific sequence is based on position weight matrices (PWMs). PWMs represent the optimal binding sequence(s) of TFs, by weighting the importance for TF binding of each of the four bases at every position in a given sequence. Such matrices have been derived for numerous TFs using chromatin immuno-precipitation sequencing (ChiP-seq) [18]. Additive models using these have been shown to accurately predict TF binding *in vitro* [19] and TFBSs with “weak” PWM-scores generally show the highest levels of individual variation [13]. Despite the potential of PWMs to calculate a score for any type of variant overlapping a TFBS, they have only been used to predict the impact of SNVs and not the one of insertions/deletions (INDELs). This is likely due to the complexities in calling and interpreting INDELs.

Two key methods, based on DNase I hypersensitive sites sequencing data (DHS-seq), emerged in the recent literature to infer the importance of variants in TFBSs [16,17]: the first uses sequencing data in the context of DNase I hypersensitivity, creating a predictive model with support vector machines (SVM) [16], while the second assesses the imbalance of sequence reads covering one or the other allele in heterozygous sites [17]. Despite the capability of the two methods to predict the consequences of variants on TF binding, both methods are limited to predicting the consequences of SNVs.

Hence a gap emerges in our capability to assess the impact of INDELs in such important regulatory regions: studies suggest that INDELs are likely to play a significant role in genome variability in health and disease [20,21], and they might be expected to introduce more severe disruptions on TFBSs than SNVs. Currently, the ability to estimate the biological role of an INDEL located outside of genes is limited to two approaches: 1) calculating the severity of the variant based on the amount of information available on the genomic region, rather than the actual change introduced to the sequence [15]; 2) correlating the INDEL to gene expression data, thereby identifying eQTLs [22].

In this investigation, we address the above mentioned gap in the understanding of how INDELs affect TF binding, by developing a method for scoring all SNVs and INDELs called in 2,504 individuals and 26 populations from the 1000 Genomes Project Phase 3 (~84 million variants) [23]. By investigating such a comprehensive resource of human genetic variation, we ensure comparable results with studies on SNVs based on experimental data and additionally provide an unprecedented window into the genome-wide landscape of INDELs and their predicted influences on TF binding. Our method provides a way to estimate the effect of variants on TF-binding that works independently of, but is comparable to, DHS-seq data for both SNVs and INDELs.

## Results

We focused our analysis on SNVs and INDELs from the 1000 Genomes Project Phase 3, combinedly accounting for about 99.4 % of the available variants in the dataset.High confidence TFBSs have been identified through the ENSEMBL API [24,25]. We have limited our analysis to TFBSs that have been mapped using a combination of ChIP-seq peaks and position weight matrices, to ensure the use of sites verified to bind TFs.

We first analysed how many of the ~84 million variants overlap known TFBSs (Table 1). Approximately 1.2 ‰ of the variants overlap at least one TFBS, 0.3 ‰ overlap at least two TFBSs while 0.1 % of the variants overlap three or more TFBSs. It is well known that TFBSs for different factors overlap, and these sites, enriched in TFBSs, are termed TF hubs [26]. It has been shown that relative levels of binding of co-occupying TFs to shared targets correlate with specific functions of these factors; furthermore TF hubs have been suggested to be particularly sensitive to genetic variation, as genetic polymorphisms have been found to destabilize TF occupancy across entire hubs [26, 27]. To ensure that most of the impact of variants on TF hubs was captured, TF binding scores were calculated for each TFBS overlapping a variant.

**Table 1:**
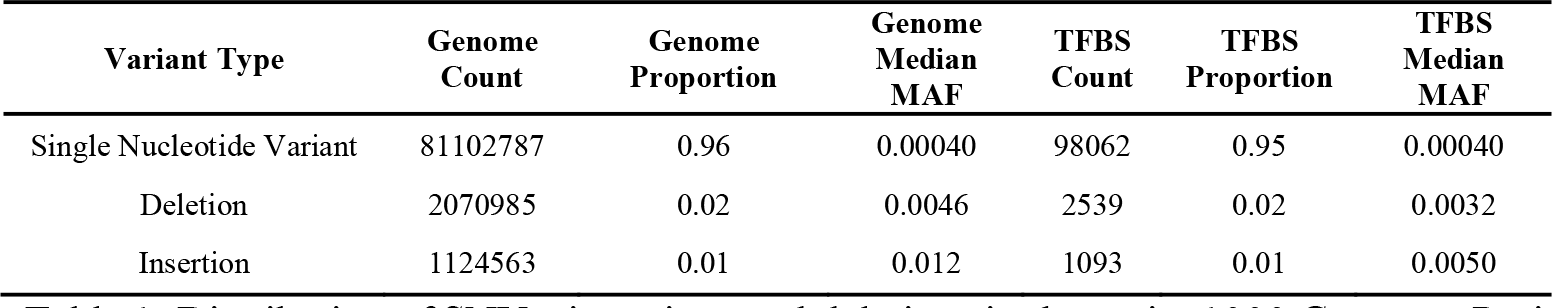
Distribution of SNVs, insertions and deletions in the entire 1000 Genomes Project Phase 3 and among variants overlapping at least one TFBS annotated in the Ensembl Regulatory Build. Genome Count: The total number of SNVs, insertions and deletions in the 1000 Genomes Project Phase 3. Genome Proportion: The proportion of SNVs, insertions and deletions of all variants in the 1000 Genomes Project Phase 3. Genome Median MAF: The median minor allele frequency (MAF) of SNVs, insertions and deletions across all populations in the 1000 Genomes Project Phase 3. TFBS Count: The number of SNVs, insertions and deletions in the 1000 Genomes Project Phase 3 that overlap at least one TFBS in the Ensembl Regulatory Build. TFBS Proportion: The proportion of SNVs, insertions and deletions of all variants in 1000 Genomes Project Phase 3 overlapping at least one TFBS. TFBS Median MAF: The median MAF of SNVs, insertions and deletions overlapping at least one TFBS.

We next compared the minor allele frequency (MAF) of all variants in the dataset to those overlapping TFBSs, by variant type: as expected, the MAF was found to be significantly lower for variants overlapping TFBSs for all three variant types (Table 1, Mann-Whitney test p-values for SNPs: 2.2×10^-16^, deletions: 2.9×10^-10^, insertions: 8.0×10^-14^). This finding is in accordance with previously mentioned studies on TFBSs indicating purifying selection [13, 28].

In order to improve our understanding of INDELs in this context, we developed a new method that uses PWMs: our method computes the effects of variants on TF binding by taking the sequence context of TFBSs into account. We introduced a sliding-window approach in our method, in order to capture new potential TFBSs originating as result of the variation. This approach makes it possible to estimate the effect of both SNVs and INDELs, overcoming most of the difficulties in evaluating the impact of deletions but also of insertions, where new sequence is introduced in the site (Figure 1). The effect is calculated as a difference between two PWM-scores that each either represents the predicted binding affinity to the TFBS of the minor allele or to that of the major allele. We thus calculated a TFBS-score that could be used to directionally estimate allelic imbalances in TF binding (Figure 1). Negative scores predict that the minor allele reduces the strength of TF binding to the site, while positive scores predict a stronger binding. TFBS-scores of zero predict no differences in TF binding between the sites containing either of the two alleles.

**Figure 1:**
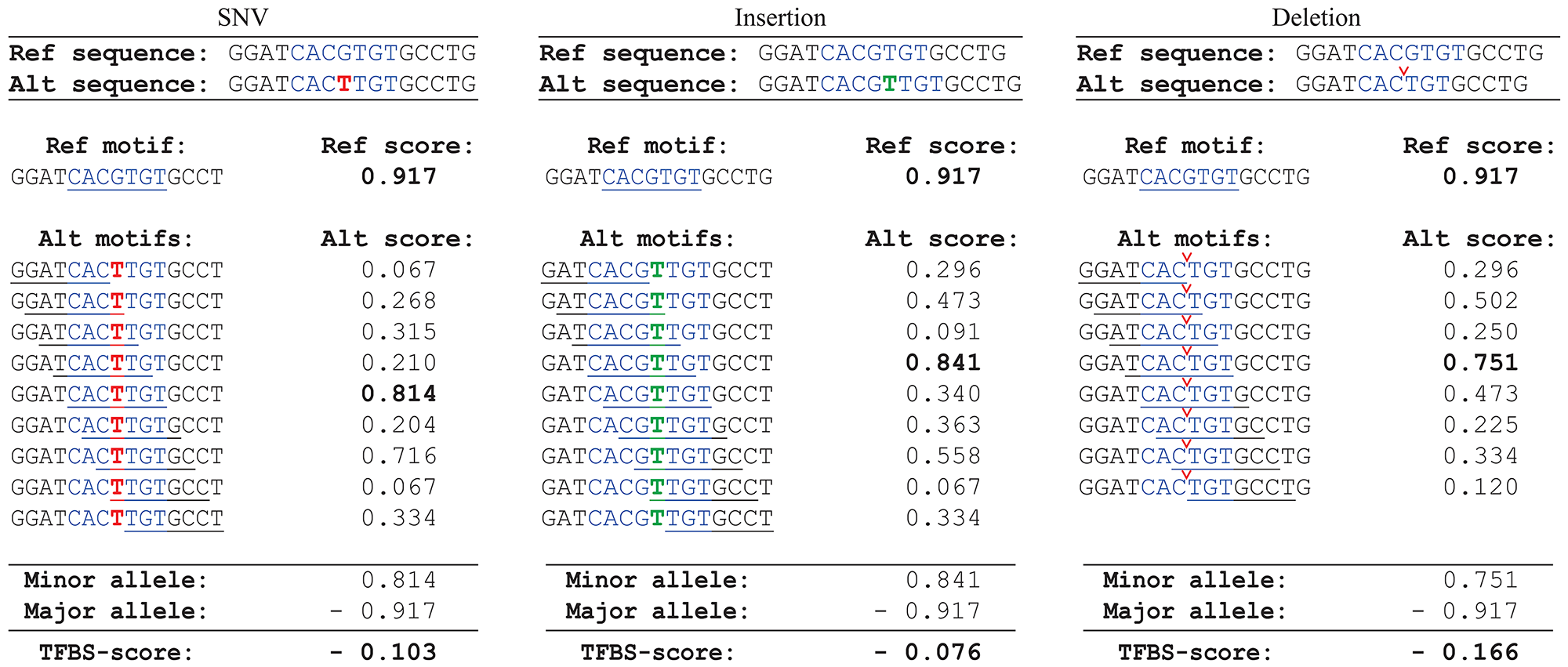
Examples of how TFBS-scores were calculated for SNVs, insertions and deletions. When estimating the PWM-score at the alternative allele, depending on the variation type, several new alternative sequences are formed that have the potential to form a new TFBSs. To ensure that these new TFBSs were captured, PWM-scores for all sequences immediately surrounding the original binding site were calculated using a sliding window approach (underlined sequences), and the surrounding sequence with the highest PWM-score was chosen as the alternative sequence (scores marked in bold). The TFBS-score was then calculated by subtracting the PWM-score at the major allele from the PWM-score at the minor allele. In all three examples above it was assumed that the major allele is the reference sequence and the minor allele the alternative sequence, though this is not always the case.

In order to thoroughly verify the consistency of our newly developed method with those investigating SNVs and based on experimental data, we used two different approaches. First, we extensively compared the TFBS-score of all our annotated SNVs with the information provided by Maurano et al. on allelic imbalanced DHS-seq reads [17]. We found that our score negatively correlates both with the percentage of reference reads across heterozygous samples (Supplementary Figure 1, Spearman's rank correlation, p-value < 2.2×10^-16^), and with their final prediction score using additional experimental data (Supplementary Figure 2, Spearman's rank correlation, p-value: < 2.2-10^-16^). Since the available experimental data contained only information about the SNVs, we called INDELs on 178 samples from the ENCODE dataset (Supplementary material 3), and applied the same method as Maurano et al., in order to measure the allelic imbalance of reads containing INDELs overlapping TFBSs. While confirming the difficulties in calling INDELs on DHS- seq data, which explain the lack of information from previous studies, we found a clear trend between our score and the allelic imbalance (Supplementary Figure 3 +4 and Supplementary Materials 3.3 for more details). We therefore concluded that our algorithm provides a way to predict the effects of INDELs on TFBSs, consistent with experimental data, but easier to compute on most datasets.

TFBS-scores were calculated for each of the 142,234 TFBSs overlapping a SNV, insertion or deletion. For each TFBS-score only a single variant was taken into account. The GERP conservation score, also available through Ensembl, was used to estimate the level of conservation for each TFBS, and for positively conserved TFBSs the TFBS-scores were found to be significantly lower than for negatively conserved TFBSs (Mann-Whitney, p-value: >2.2×10^-16^). The distributions of TFBS-scores differed significantly between the three variant types (Figure 2; Kruskal-Wallis, p-value: >2.2×10^-16^). All three distributions were found to have the highest densities around zero, predicting that the majority of variants have little or no effect on the TF binding affinity. This finding is in agreement with experimental studies suggesting that only a minority of TFBSs containing heterozygous variants show significant allele-specific TF binding [29,30]. This trend was surprisingly the most pronounced for insertions, whereas deletions, as expected, on average had the most severe effects. The three distributions were compared pairwise to estimate their similarity, and the distribution of TFBS-scores for insertions was found to have the most distinct profile (Mann× Whitney, p-values for SNVs vs Insertions: 1.58×10^-98^, SNVs vs deletions: 8.39×10^-6^, insertions vs deletions: 8.7×10^-62^).

**Figure 2:**
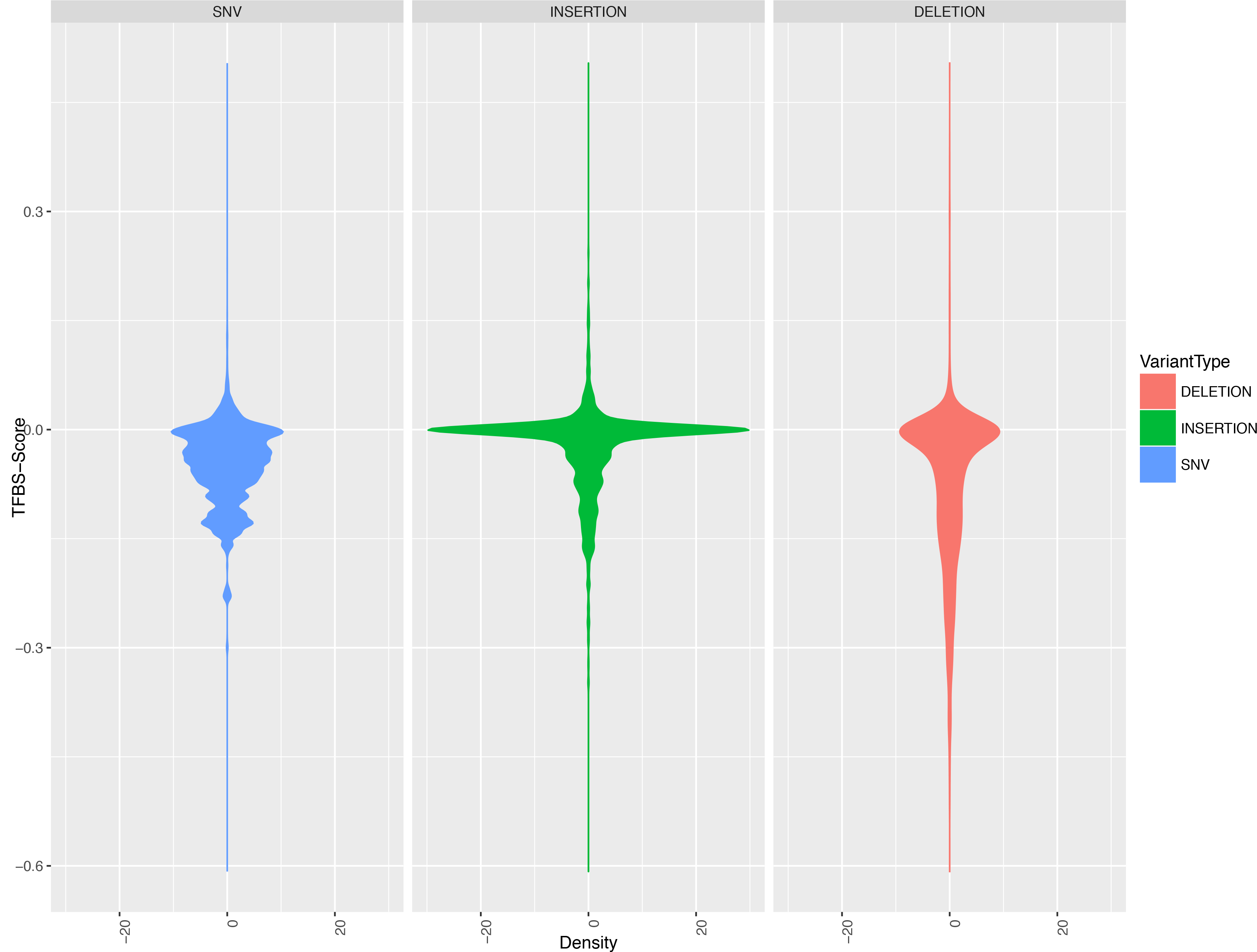
Violin plot of the TFBS-score distributions for each of the 142,234 TFBSs overlapping a SNV, insertion or deletion. The x-axis indicates the TFBS-score density mirrored around zero. The y-axis indicates the TFBS-score.

Next we investigated how the position of variants within TFBSs relates to the TFBS× score (Figure 3A). For all three variant types on average the TFBS-scores were the closest to zero in the start and in the end of TFBSs, whereas the scores were the most negative in the centre of the sites, more clearly for INDELs. This trend inversely correlated with the average information content across the 63 TFBSs investigated, for which the average information content was found to be the highest in the centre of the sites and the lowest at the edges (Supplementary Figure 7). As previously seen (Figure 2), deletions on average resulted in the most negative scores, whereas insertions resulted in the least negative. Next we calculated the average MAF for the variants across the TFBSs (Figure 3B). The overall MAF of variants seemed to be almost evenly distributed across the TFBSs for all three variant types. We therefore investigated the relationship between the MAF and the TFBS-score: for all three variant types the least common variants were on average found to have the most severe TFBS-scores (Figure 4), and a significant correlation between the two were found for each of the three variant types (Spearman's rank correlation, p-values for SNVs: p-value < 2.2×10^-16^, Insertions: 8.46×10^-13^ and Deletions: < 2.2×10^-16^).

**Figure 3:**
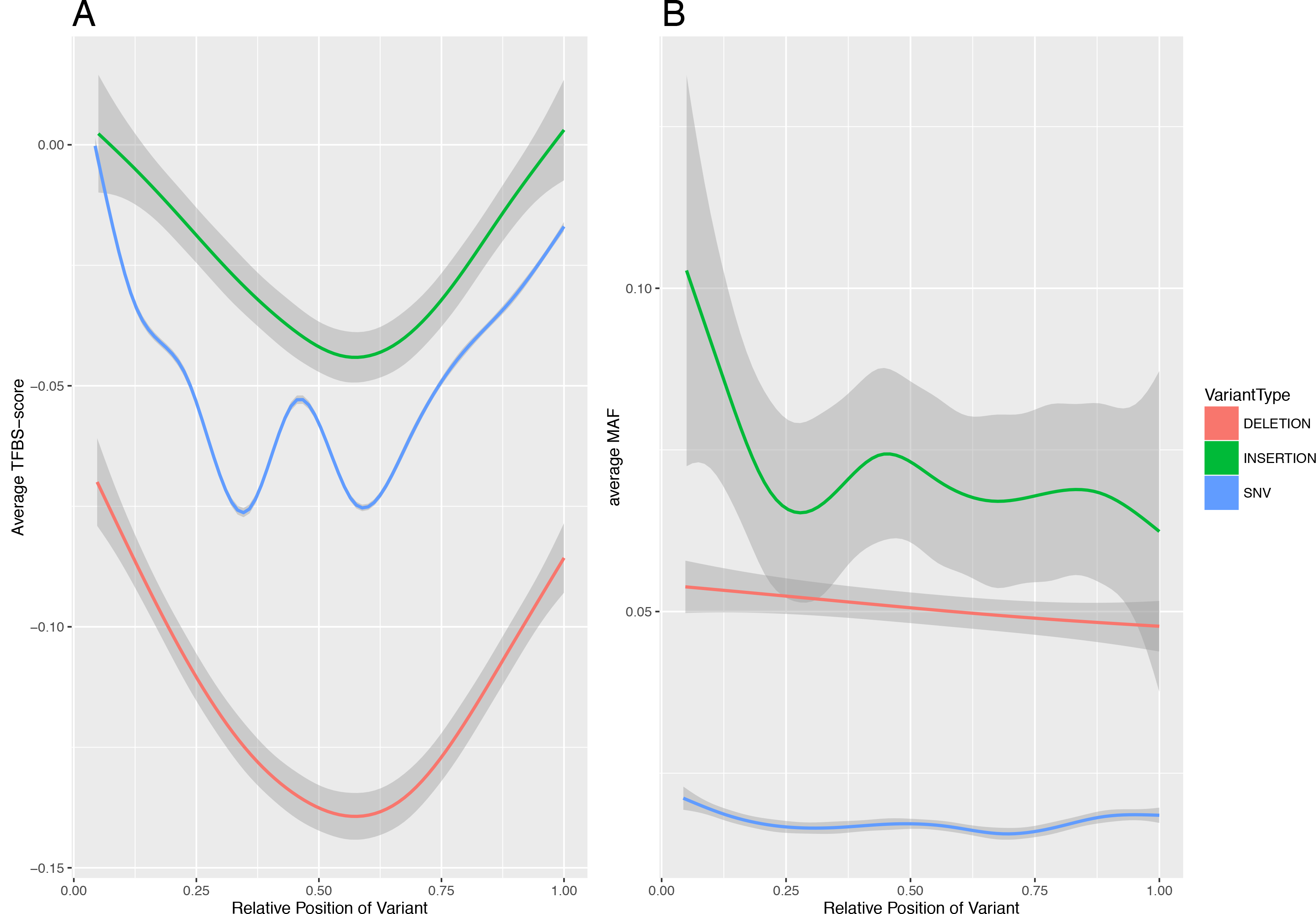
The relationship between variant position and both TFBS-scores (A) and MAF (B). Deletions were represented as SNVs at each position in the site.

**Figure 4:**
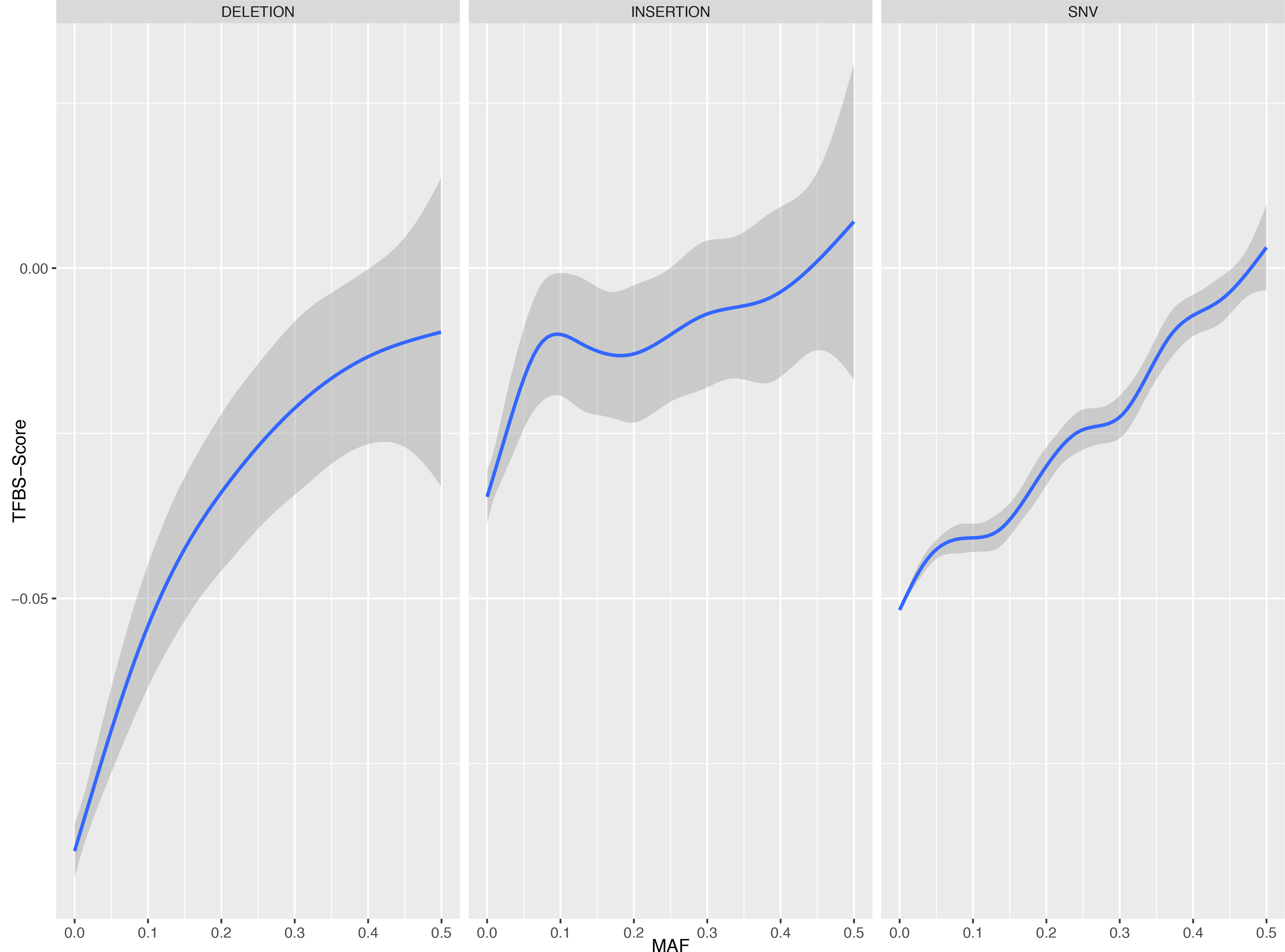
TFBS-score vs MAF per variant type.

To understand the clinical relevance of the TFBS-scores, the overall TFBS-score distribution was divided into four categories based on the two major local minima of the distribution, and an interval around zero was identified using the local minimum closest to zero (Figure 5A; from left to right): Category 1) Variants with very negative TFBS-scores, Category 2) with moderately negative scores, Category 3) with scores around zero, Category 4) and variants with positive scores. Each of the four categories was tested for enrichment in clinically relevant variants according to the categories described in the ClinVar database [31].

Category 1, with the most negative scores, was the one most significantly enriched in pathogenic variants (Figure 5B; Category 1, p-value: 4.1×10^-6^), suggesting that variants with larger magnitude in reduction of TF binding affinity are more likely to be pathogenic. Category 2, with negative TFBS-scores, is significantly enriched in both pathogenic (p-value: 4.8×10^-3^) and benign (p-value: 2.1×10^-10^) variants (Figure 5B; Category 2). In category 3 there was significant enrichment for more variant classes: pathogenic (p-value: 1.1×10^-2^), benign (p-value: 1.22×10^-3^) and protective (p-value: 5.4×10^-7^) variants (Figure 5B; Category 3). Finally, category 4 was significantly enriched only in benign variants (Figure 5B; Category 4, p-value: 7.1×10^-2^).

**Figure 5:**
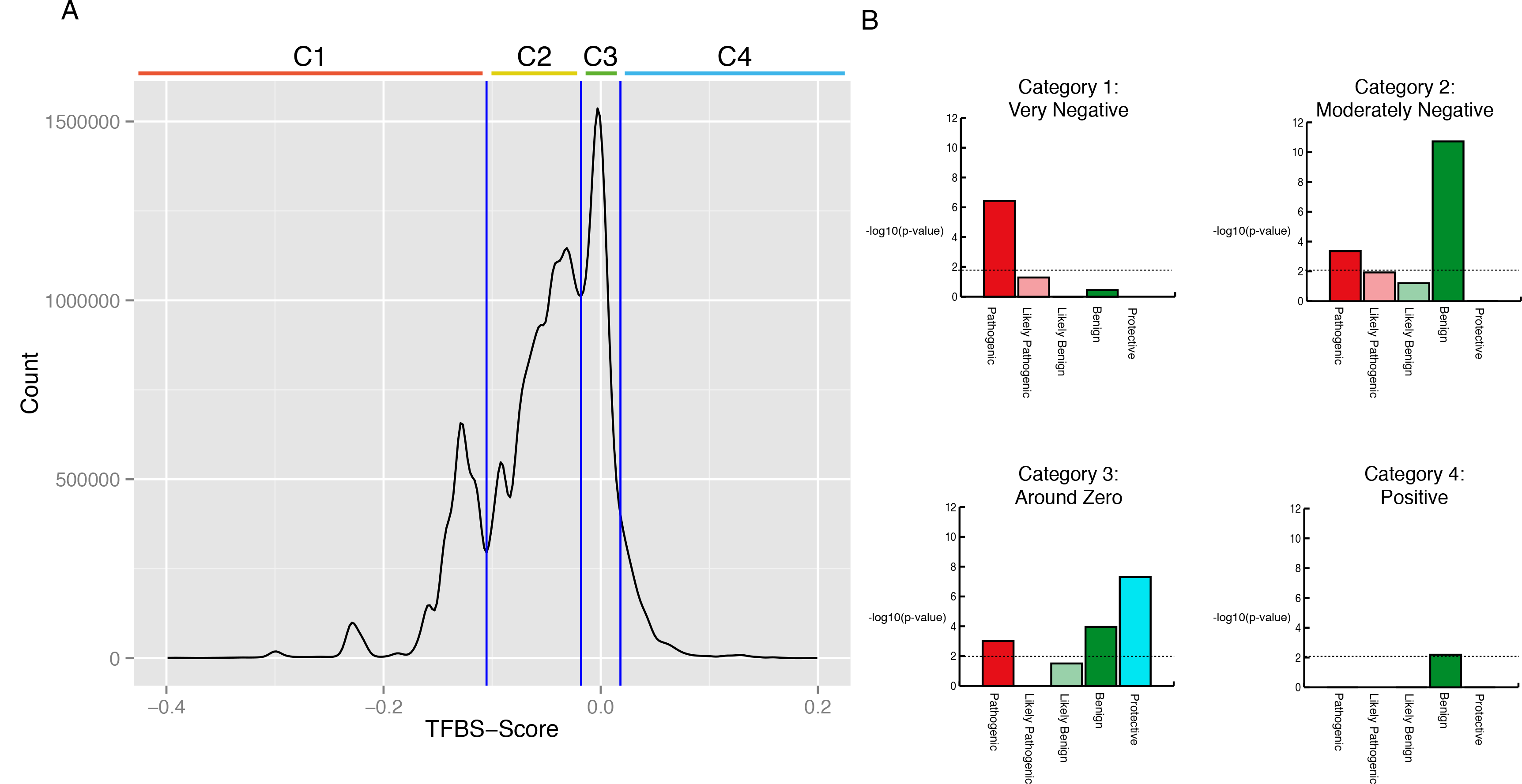
The variants were divided into four categories based on their TFBS-score distributions, and each category was tested for enrichment in clinically relevant variants from the ClinVar database. A) The overall TFBS-score distribution divided by three vertical blue lines into four categories based on the two major local minima of the distribution, and an interval around zero was identified using the local minimum closest to zero. From left to right the categories are: Category 1 (C1): Very Negative, Category 2 (C2): Moderately Negative, Category 3 (C3): Around Zero, Category 4 (C4): Positive. B) The results from the enrichment analysis with-log10 of the p-values on the y-axis and the ClinVar categories on the x-axis. The dotted lines represent a significance level of 0.05 Bonferroni adjusted.

To better understand the differences in variation between the different TFBSs we calculated the average TFBS-score for each (Figure 6). The score profiles of the different TFBSs were quite diverse: interestingly, the one with the lowest average score binds GATA2, while the one with the highest average scores binds CTCF. Both TFs have important roles, and have been associated to various pathologies.

**Figure 6:**
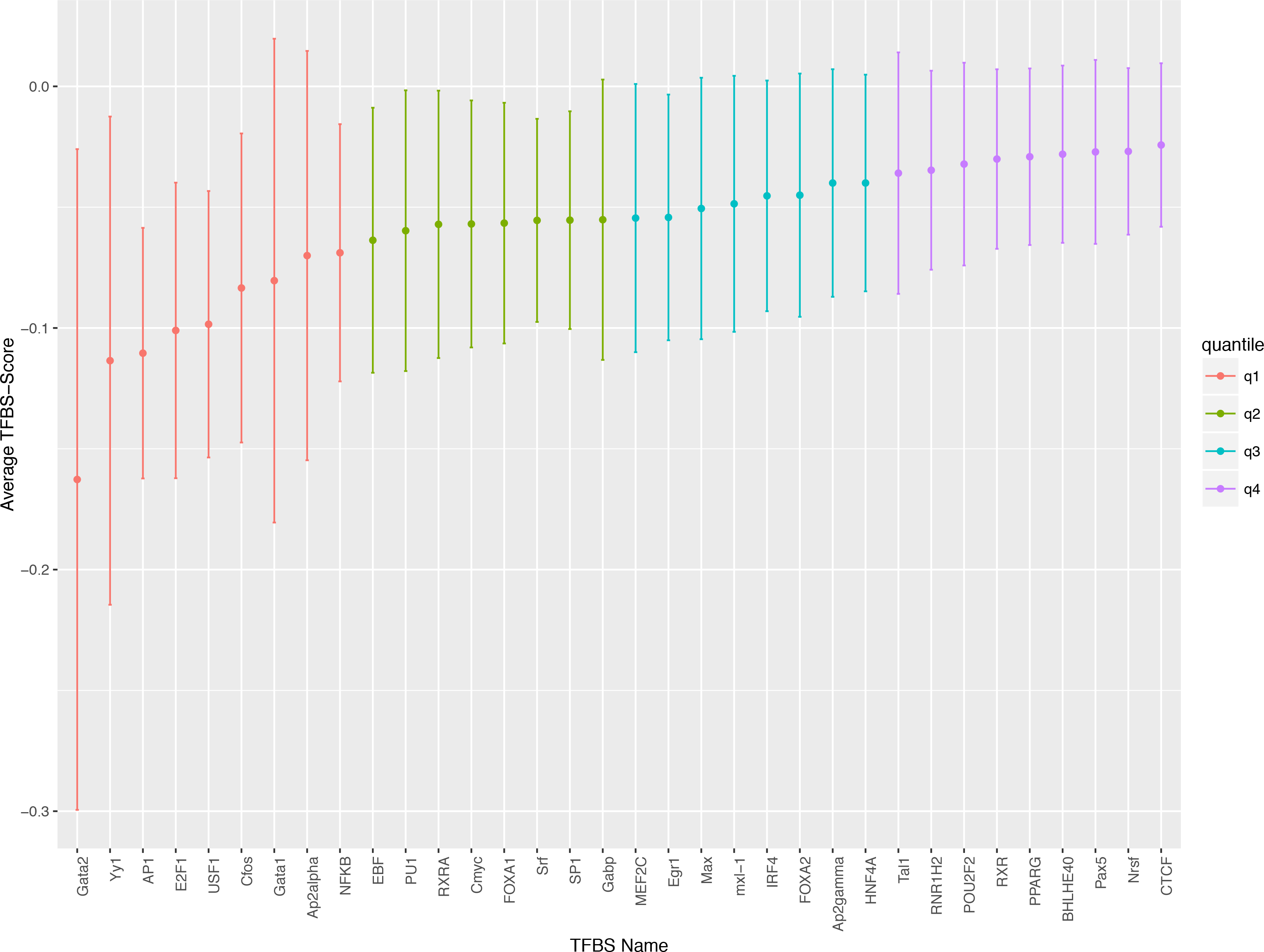
Average TFBS-scores for each TFBS with the whiskers indicating the standard deviation.

## Discussion

In this paper we address a significant gap in the literature, as well as a shortcoming of currently available bioinformatic tools to investigate transcription factor binding sites. The lack of in-depth information on the role of insertions/deletions on TFBSs is a key limitation in our understanding of genetic variants located in non-coding regions. The reason for this gap is mainly due to difficulties in calling INDELs from Chip-Seq or DHS-Seq data: sequence occupancy experiments still rely on short reads for identifying TF footprints [32], and short reads by definition provide a low-quality material to call insertions/deletions. Therefore, bioinformatic tools that can be used on any genotype dataset, without the need of additional experiments, can be a great alternative to estimate the effects of variants in TFBSs. We validated our approach, by checking the correlation of our annotation with those available for SNVs calculated using other methods [17]. Additionally, we have applied the same thorough approach of previous studies [17] in order to measure the allelic imbalance at sites where insertion/deletions overlap TFBSs, by calling insertion/deletions from raw data of a large number of individuals available in ENCODE, and correlated our results with DHS-Seq data also for INDELs. We can thus conclude that our method provides a reliable measure to investigate the effect of genetic variation on TF binding across the genome.

We used this newly developed approach to extensively characterise all SNVs and INDELs from 1000 Genome phase 3 and provide a comprehensive description of all known INDELs located in TFBSs. The picture we provide with this approach confirms some of the major findings for single nucleotide variants in these regions: the average MAF of INDELs overlapping TFBSs has been found significantly lower than the average MAF of all INDELs in the dataset. Similarly, the TFBS-score has been found to be significantly lower for highly conserved TFBSs compared to less conserved TFBSs, i.e. the variability of the sequence being consistent with a locus under evolutionary constraints. It is also known that flanking regions of TFBSs harbour higher variation than the central regions, where the information content is higher because the sequence is likely more important for the protein recognition and binding. We confirm these characteristics also for INDELs by using our score: insertions/deletions in the edge regions of binding sites have higher scores than those located in the centre of TFBSs, indicating that INDELs in edge regions of TFBSs are less disruptive to binding while INDELs at the centre of TFBSs are more likely to reduce the strength of TF binding.

Our findings for deletions are in line with the expectation that these variants introduce more severe disruptions of the binding sequences than SNVs; however, insertions seem on average to display less severe consequences than SNVs. We might speculate that, while deletions ablate part of the binding site, insertions introduce an additional sequence, by splitting but not removing the original binding sequences. This different action, and potential of creating alternative binding sequences, as we verify with our sliding window approach (see methods and Figure 1), might account for the less severe consequences we observed in the binding affinities of TFs.

A major reason for devoting time to understanding better the biological consequences of genetic variants is the need for effective tools to interpret the increasing amount of sequence data available in clinical settings and in precision medicine initiatives. For this reason we analysed the distribution of the scores calculated with our method, using the information available in ClinVar, a progressively more important resource for the annotation of variants with potential for clinical relevance. In the current version of the database, the variants are flagged as pathogenic, likely pathogenic, benign, likely benign and protective, depending on the amount of support for the pathological consequences of each genetic variant [31]. We tested for enrichment of these variant groups in four categories of TFBS-scores and our results confirmed the hypothesis that variants with the most negative scores, i.e. those most severely reducing binding affinity, are the most enriched in pathogenic variants. This is relative straightforward to interpret: those variants that severely disrupt binding sites, especially when those are evolutionary conserved, are most likely to produce a negative effect and therefore be pathogenic.

On the other hand, both categories with moderately negative (Category 2) and positive scores (Category 4) show a positive enrichment in benign variants (Figure 5). We might speculate that in both cases the effect of the changes to the binding affinity does not have sufficient magnitude to be either damaging or protective: indeed, variants in category 2 are only slightly reducing the binding affinity, while variants in category 4 are only slightly increasing the binding affinity.

More surprisingly, variants with scores around zero (Category 3) are the only group significantly enriched in protective variants, but also a group enriched both in pathogenic and benign variants, to a lower extent. These mixed enrichment results for variants with scores around zero are less straightforward to decipher. A closer look at the variants in this group reveals the presence of quite heterogeneous variation, and the clinical importance seems to be independent from the effect on TF binding. In this category we found, for example, a SNV (rs2302615) which is classified as “protective”: it overlaps three TF binding sequences and its TFBS-scores range between -0.0012 and 0.0151, indicating no large effect on TF binding; however, rs2302615 is associated with reduction in adenoma recurrence when the subjects are treated with aspirin [33], which is the reason for its classification. On the other hand, in this category we also find variants like rs72553883, which also has no influence on TF binding (TFBS-score = 0.0153), but is reported to be a missense variant causing severe impairment in B-cells function, and therefore is associated to common variable immunodeficiency [34]. The variant is annotated as “pathogenic”, but its biological consequence is clearly related to the amino acid change, rather than to the effect on the overlapping TFBS. The mixed enrichment results we observed for this category seem therefore due to reasons independent of the effect on the TF binding.

The scoring method we present here also allows us to look at transcription factors from a different perspective: we show that we can describe a characteristic profile for each TF, based on the scores of the variants affecting its binding sites across the whole genome.Interestingly, if we look at those profiles, we find that CTCF is the TFBS with the highest average score, and GATA2 is the one with the lowest. They both play a pivotal role in gene regulation. CTCF is a transcription factor that once bound to DNA, can function as a transcriptional insulator, repressor or activator. A number of mutations in the CTCF gene have been associated with mental retardation [35]. GATA2 is a transcription factor expressed in hematopoietic progenitors, including early erythroid cells, mast cells and megakaryocytes. Mutations in the GATA2 gene have been associated with both *Immunodeficiency 21* and *Chronic Myeloid Leukemia* [36,37].

## Conclusions

With this work we introduce a way to estimate the effects of variants on TF binding, independently of the availability of additional experimental data for the genetic dataset to be analysed. Our method handles both SNVs and INDELs, thus filling an important gap in the interpretation of the effect of insertion/deletions in TFBSs. The enrichment analysis, finally, shows how our score can be used in order to further prioritise variants in sequencing studies.

## Materials and methods

### Data sources

Human variation data, including allele frequencies, are from the 1000 Genomes Project Phase 3 release based on the 20130502 sequence freeze and alignments with file date 18^th^ of February 2015 [23]. TFBSs, PWMs and genetic annotations were detected querying the Ensembl API version 75, assembly GRCh37 [25] using a custom script. The following four databases were installed locally for the queries: homo_sapiens_core_75_37, homo_sapiens_funcgen_75_37, homo_sapiens_variation_75_37 and ensembl_compara_75. Variants annotated with clinically relevance were retrieved from the variant summary of the ClinVar database May 25^th^ [31]. Phenotypes and ontologies were retrieved using the R package biomaRt [38]. Analyses were performed using BASH and R.

### Motif selection for analysis

The motifs with overlapping variants were detected querying the Ensembl functional genomics database. In the database putative TFBSs live up to the two criteria; there has to be both ChIP-seq data and publicly available PWMs available. For each variant all overlapping MotifFeatures were detected. MotifFeatures are objects representing the genomic location of a sequence motif. If the same motif was detected twice for a variant, only the motif with the most negative TFBS-score was selected. For each MotifFeature information was available on what TF bound the motif and what PWM was associated with it.

### Transcription factor binding site scores

For each TFBS with an overlapping variant a TFBS-score was calculated. The score predicts the difference in TF binding affinity between the TFBS where the major allele is present and the TFBS with the minor allele. The TFBS-score was calculated in two steps using PWMs available through the Ensembl API. First, PWM-scores, predicting the individual TFBS binding affinities, were calculated for the major and the minor allele.

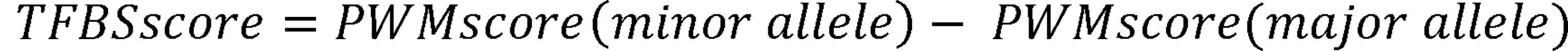

The PWM-scores were calculated using a method available through the Ensembl API called “Bio::EnsEMBL::Funcgen::BindingMatrix::relative_affinity()”. This method calculates the binding affinity of a given sequence relative to the optimal site for the matrix, considering a random background, where the probability of each of the four bases was considered to be equal at each position. The method returns scores between 0 and 1, where 1 means that the sequence the score is calculated for is the optimal binding sequence. Depending on the variation type (SNVs, insertions or deletions), several new sequences were formed at the minor allele with the potential to form a new TFBS. To account for this issue, PWM-scores for all sequences immediately surrounding the original binding site were calculated for SNVs, insertions and deletions. For insertions a total of TFBS length + insertion length + 1 sequences were scanned for a new TFBS, while for deletions a total of TFBS length + 1 sequences were scanned and for SNVs a total of TFBS length + 2 sequences. The surrounding sequence with the highest PWM score was then chosen to represent TF binding at the minor allele.

### Filtering of ClinVar data

ClinVar contains reports of the relationships among human variations and phenotypes including supporting evidence. We used the variants with rs-numbers together with their genomic position relative to the GRCh37 reference. Variants annotated both with benign and pathogenic effects were flagged as conflicting data. All variants annotated with uncertain significance were also flagged as conflicting data no matter the additional variant annotations. In order to be conservative variants that were flagged as both being likely benign and benign were flagged as likely benign, and the same approach was used to flag variants that were both annotated as likely pathogenic and pathogenic. Variants with only one annotation were flagged accordingly.

### Enrichment analysis

We used a hypergeometric test to identify which groups of clinically relevant variants, as classified in ClinVar, were over-represented in the four categories of variants we identified in our TFBS-score distribution. The variant population used for the tests included all variants from the 1000 Genomes Project Phase 3 potentially overlapping variants annotated in the ClinVar database. That is all SNVs, insertions and deletions in the 1000 Genomes phase 3 with rs-numbers. The variants analysed for enrichment included all SNVs, insertions and deletions annotated in the ClinVar database with rs-numbers.

### Availability of data and materials

Script used to annotated The 1000 Genomes Project Phase 3:

> Project name: TFBS Annotator
>
> Project home page: github.com/esbeneickhardt/TFBSAnnotator
>
> Operating system(s): Platform independent
>
> Programming language: Perl
>
> Other requirements: ENSEMBL API version 75
>
> License: MIT modified
>
> Any restrictions to use by non-academics: licence needed

Dataset with all TFBSs in the ENSEMBL databases that overlap a variant in the 1000 Genomes Phase 3:

> github.com/esbeneickhardt/TFBSAnnotator/tree/master/Data/TFBSDataset.txt.zip.

Dataset with all TFBSs in the ENSEMBL databases that overlap a variant called in the public DHS data (GSE26328).

> github.com/esbeneickhardt/TFBSAnnotator/tree/master/Data/DHSDataset.txt

## Abbreviations

ChiP: chromatin immunoprecipitation
DHS: DNase I hypersensitive site
eQTL: expression quantitative trait loci
GWAS: genome-wide association study
INDEL: insertion/deletion
MAF: minor allele frequency
PWM: position weight matrix
SNV: single nucleotide variant
SVM: support vector machines
TF: transcription factor
TFBS: transcription factor binding site

## Authors' contributions

EE co-designed the study, wrote the motif discovery and annotation script, conducted the analysis and wrote the manuscript. FL supervised, conceptualized and co-designed the study, contributed substantially to all aspects thereof, and contributed to writing the manuscript. TDA overlooked evolutionary aspects of the study, and contributed to the interpretation of the results. JG overlooked statistical aspects of the study. AB provided funding for the study and supervised the work. All authors have read and approved the final manuscript.

## Acknowledgements

The authors wish to thank Jonatan Pallesen for valuable advice and input on how to bioinformatically and methodically address various challenges, and to thank all members of the B0rglum lab for helpful discussions. This work was funded by the iPSYCH - Lundbeck Foundation Initiative for Integrative Psychiatric Research.

